# BIN2 phosphorylates the Thr280 of CO to restrict its function in promoting *Arabidopsis* flowering

**DOI:** 10.1101/2022.04.20.488915

**Authors:** Lan Ju, Huixue Dong, Ruizhen Yang, Yexing Jing, Yunwei Zhang, Liangyu Liu, Yingfang Zhu, Kun-Ming Chen, Jiaqiang Sun

**Affiliations:** National Key Facility for Crop Gene Resources and Genetic Improvement, Institute of Crop Sciences, Chinese Academy of Agricultural Sciences, Beijing 100081, China; Sorghum Institute of Shanxi Agricultural University, Jinzhong 030600, China; State Key Laboratory of Crop Stress Biology in Arid Areas, College of Life Sciences, Northwest A&F University, Yangling 712100, Shaanxi, China; State Key Laboratory of Crop Stress Adaptation and Improvement, School of Life Sciences, Henan University, Kaifeng 475001, China; Beijing Key Laboratory of Plant Gene Resources and Biotechnology for Carbon Reduction and Environmental Improvement, and College of Life Sciences, Capital Normal University, Beijing 100048, China

**Keywords:** BIN2, CO, function, phosphorylation, flowering, *Arabidopsis*

## Abstract

CONSTANS (CO) is a central regulator of floral initiation in response to photoperiod. In this study, we show that the GSK3 kinase BIN2 physically interacts with CO and the gain-of-function mutant *bin2-1* displays late flowering phenotype through down-regulation of *FT* transcription. Genetic analyses show that BIN2 genetically acts upstream of CO in regulating flowering time. Further, we illustrate that BIN2 phosphorylates the Thr280 residue of CO. Importantly, the BIN2 phosphorylation of Thr280 residue restricts the function of CO in promoting flowering. Moreover, we reveal that the N-terminal part of CO harboring the B-Box domain mediates the interaction of both CO-CO and BIN2-CO. We find that BIN2 inhibits the formation of CO dimer/oligomer. Taken together, this study reveals that BIN2 regulates flowering time through phosphorylating the Thr280 of CO and inhibiting the CO-CO interaction in *Arabidopsis*.

**Highlight:** BIN2 regulates flowering time through phosphorylating the Thr280 of CO in *Arabidopsis*.

## Introduction

Flowering is a transition from the vegetative to the reproductive phase in the life cycle of flowering plants, which is controlled by several pathways, including photoperiod, autonomous, vernalization, age and gibberellin pathways (Andrés and Coupland, 2012). Among these pathways, photoperiod pathway plays a most important role in controlling *Arabidopsis thaliana* flowering time (Komeda, 2004). CONSTANS (CO) and FLOWERING LOCUS T (FT) are the central regulators of floral initiation in response to photoperiod (Putterill et al., 1995; Kobayashi et al., 1999). CO is a B-Box-type zinc finger transcription factor that promotes flowering by directly activating *FT* mRNA expression through binding to the two CO-responsive elements (CORE1 and CORE2) in the *FT* promoter (Onouchi et al., 2000; Samach et al., 2000; Tiwari et al., 2010; Lv et al., 2021).

To date, several components have been identified to precisely regulate the diurnal transcription of *CO* in *Arabidopsis*. For example, CYCLING DOF FACTORs (CDFs) repress flowering by down-regulating *CO* transcription in the leaves (Sawa et al., 2007). The bHLH transcription factors FLOWERING BHLHs (FBHs) directly activate *CO* transcription to promote flowering (Ito et al., 2012). The TEOSINTE BRANCHED/CYCLOIDEA/PCF 4 (TCP4) transcription factor directly interacts with FBHs to synergistically activate *CO* transcription and promote flowering (Liu et al., 2017). A recent study reported that the histone demethylase JMJ28 regulates *CO* by interacting with FBH transcription factors to promote flowering (Hung et al., 2021). At the posttranscriptional level, the activity and stability of CO are also regulated. For example, NUCLEAR FACTOR Y (NF-Y) transcription factors physically interact with CO to activate *FT* expression in regulating flowering; TARGET OF EAT1 (TOE1) interacts with CO and inhibits its activity (Kumimoto et al., 2010; Zhang et al., 2015). In addition, CO can be ubiquitinated by CONSTITUTIVE PHOTOMORPHOGENIC1 (COP1) and degraded by 26S proteasome-dependent proteolysis (Jang et al., 2008; Liu et al., 2008). Interestingly, a previous study reported that CO can be phosphorylated, which contributes to the photoperiodic flowering response by facilitating the rate of CO degradation via the COP1 ubiquitin ligase (Sarid-Krebs et al., 2015).

Brassinosteroids (BRs) play a vital role in controlling floral transition. Previous studies have shown that BR biosynthetic and signaling mutants, including *de-etiolated 2 (det2), constitutive photomorphogenesis dwarfism (cpd), brassinosteroid insensitive 1-1* (*bri1-1*) and *bin2-1*, display late-flowering phenotype (Li et al, 2002; Clouse, 2008; Li et al., 2010). However, the fundamental mechanism of BR-regulated flowering remains obscure. Up to now, BR signal transduction cascade from the cell surface receptor kinase BRI1 to the BRASSINAZOLE RESISTANT 1 (BZR1) family transcription factors has been well clarified (Wang et al., 2012; Chaiwanon et al., 2016). In the presence of BR, BRs bind to and activate BRI1, leading to inhibition of the GSK3-like kinase BRASSINOSTEROID INSENSITIVE 2 (BIN2) (Yan et al., 2009; Kim and Wang, 2010). Upon BIN2 inactivation by upstream BR signaling, BZR1 family transcription factors are dephosphorylated and accumulated in nuclear to regulate BR-responsive gene expression (He et al., 2002; Wang et al., 2002; Yin et al., 2002).

In this study, we demonstrate that BIN2 physically interacts with CO to phosphorylate its Thr280 residue, causing late flowering. On the other hand, we find that BIN2 prevents the formation of CO dimer/oligomer. Taken together, we uncover a regulatory module BIN2-CO in coordinating flowering time in *Arabidopsis*.

## Materials and Methods

### Plant materials and growth conditions

*Arabidopsis thaliana* plants used in this study are in Col-0 ecotype background. The following mutants and transgenic lines used in the study have been previously described: *bri1-116* (Friedrichsen et al., 2000), *bin2-1* (Li and Nam, 2002), *det2-9* (Lv et al., 2018), *co-9* (Balasubramanian et al., 2006), *proCO:HA:CO* (Hayama et al., 2017; Cheng et al., 2020)*, 35S:Flag-BIN2* (He et al., 2019). Seeds were grown on half-strength Murashige and Skoog medium and stratified at 4°C for 2 d, then grown in long-day (16 h light/8 h dark) conditions at 22°C. Time-course analyses were performed on 14-d-old seedlings. *Nicotiana benthamiana* plants were grown under LD (16 h light/8 h dark) conditions at 24 □.

### Analyses of flowering time phenotype

Analyses of flowering time were performed as previously described (Liu et al., 2017). Flowering time was scored as the number of days from germination to the first appearance of buds at the apex (days to bolting). More than 15 plants were counted and averaged for each measurement.

### DNA constructs

For Gateway cloning, all the gene sequences were cloned into the entry vector *pQBV3* (Gateway) and subsequently introduced into certain destination vectors following the Gateway technology (Invitrogen) (He et al., 2019). For ligase-independent ligation, the ligation free cloning mastermix (abm) was used following the application handbook. For generation the mutant forms of *BIN2* and *CO*, their coding sequences were first cloned into the *pQBV3* vector, and then the point mutations were introduced by the specifically designed primers as previously described (Zheng et al., 2004), following the PCR amplification program: preheating at 94°C for 3 min, 16 cycles of 94°C for 1 min, 55°C for 1 min and 68°C for 4 min, finally, added DpnI and incubated at 37°C for 4 h. All details of DNA constructs and the primers used in this study are listed in Supplemental Table 1.

### Generation of transgenic/hybrid plants

For the generation of *35S:YFP-CO* transgenic line, the full-length *CO* coding sequence was cloned into *p35S-YFP* vector to generate the *35S:YFP-CO* construct. The *35S:YFP-CO* construct was then transformed into the *Agrobacterium* strain *GV3101* and introduced into Col-0 by the floral-dip method (Clough and Bent, 1998). For the generation of *pCO:Flag-CO* and *pCO:Flag-CO^T280A^* transgenic lines, the fragment containing the *CO* promoter and the *CO/CO^T280A^* coding region were cloned into the *pCAMBIA1301* vector to generate the *pCO:Flag-CO* and *pCO:Flag-CO^T280A^* constructs, respectively. Then the *pCO:Flag-CO* and *pCO:Flag-CO^T280A^* constructs were transformed into the *Agrobacterium* strain *GV3101* and introduced into the *co-9* mutant background by the floral-dip method.

The *35S:YFP-CO/bin2-1* line was generated through genetic crossing between the *35S:YFP-CO* and *bin2-1* lines. The *35S:YFP-CO/35S:Flag-BIN2* double transgenic plants were generated by genetic crossing between the *35S:YFP-CO* and *35S:Flag-BIN2* transgenic lines.

### RNA extraction and quantitative real-time PCR

Total RNAs were extracted using a plant total RNA extraction kit (Zoman). Total RNA concentration was measured by Nanodrop 2000 (Thermo Fisher Scientific NanoDrop). About 2 μg of total RNA was used for reverse-transcribing to cDNA by abm reverse transcriptase kit. The cDNA was diluted to 100 μL with water in a 1:5 ratio, 2μL diluted cDNA was used for qRT-PCR. And qRT-PCR was performed with SYBR Premix Ex Taq (TaKaRa) on Light Cycler 96p (Roche). *ACTIN7* was used as an internal control. Experiments were performed independently three times with similar results. All the primers used for qRT-PCR are listed in Supplemental Table 1.

### LCI assays

The LCI assays were performed in *N.benthamiana* leaves as described previously (He et al., 2019, Yang et al., 2021). Briefly, the full-length or truncated forms of the genes were cloned into the *pCAMBIA1300-nLUC* and *pCAMBIA1300-cLUC* vectors. Different constructs were transformed into the *Agrobacterium* strain *GV3101* and then co-infiltrated into *N. benthamiana* leaves. Then plants were incubated for 12 h under dark and 36 h under light, and the LUC activities were analyzed using NightSHADE LB985 (Berthold). For each protein-protein interaction assay, at least 5 independent *N. benthamiana* leaves were infiltrated and analyzed, and 3 independent biological replications were performed for each assay (Sun et al., 2013).

### Yeast two-hybrid assays

For yeast two-hybrid (Y2H) analysis, the full-length or truncated forms of the genes were cloned into vectors *pGADT7* or *pGBKT7*, respectively. Different constructs were transformed into *AH109* yeast cells, and grown on agar plates with SD–L/W (synthetic dextrose medium lacking Leu and Trp). Further, the yeast cells were screened on SD–L/W/H medium (synthetic dextrose medium lacking Leu, Trp and His). Plates were kept at 30□ for 3 to 4 days. The empty vectors *pGADT7* and *pGBKT7* were used as negative controls.

### Co-IP assays

For the co-immunoprecipitation (Co-IP) assays, the *35S:YFP-CO* and *35S:YFP-CO/35S:Flag-BIN2* transgenic plants were used for detecting the interaction between CO and BIN2. Samples were harvested at ZT16 and then grounded to fine powder in liquid nitrogen. The total proteins were extracted by the lysis buffer (50 mM Tris-HCl at pH 7.5, 150 mM NaCl, 5 mM EDTA at pH 8.0, 0.1% Triton X-100, 0.2% NP-40) with freshly added PMSF (phenylmethylsulphonyl fluoride, 10 mM), Roche protease inhibitor cocktail (Roche, 11873580001) and MG132 (20 μM). After centrifugating at 4°C for 20 min, Extracts were mixed with Protein-G (Invitrogen, 00621751) for 1 h to reduce nonspecific immunoglobulin binding, and then incubated with anti-GFP magnetic beads (MBL, M047-8) overnight at 4°C. The precipitated samples were washed at least five times using the lysis buffer and then added 2X SDS protein loading buffer boiling for 5 min. The beads were detected by anti-GFP (1:2000; Roche, 11814460001) and anti-Flag (1:5000; abmart, M20026) antibodies.

### *In vitro* pull-down assays

The full-length coding sequences of CO and BIN2 were cloned into the *pGEX4T-1* and *pMAL2X-1* vectors to generate GST-CO and MBP-BIN2, respectively. These fusion proteins were expressed and purified from *E. coli* strain *Transetta-DE3* (Transgen biotech, CD801). The pull-down assays were performed as described previously (Yang et al., 2021). For the assay, prewashed amylose resin in the column buffer (20 mM Tris [pH 8.0], mM NaCl, 1 mM EDTA) was used to pull down protein complexes. The mixture with proteins and amylose resin was incubated overnight at 4 °C, then washed for 5 times with wash buffer, and added 2X SDS protein loading buffer boiling for 5 min. Eluted proteins were analyzed by immunoblotting using anti-MBP (1:3000; CWBIO, CW0288) and anti-GST (1:3000; CWBIO, CW0144) antibodies.

### Protein extraction and immunoblotting

For protein level analysis, every 100 mg of *Arabidopsis* seedlings or *N. benthamiana* leaves added 200 μl protein extraction buffers. The extracted buffer used for extracting plants is described previously (He et al., 2019, Yang et al., 2021). The supernatant was separated on SDS-PAGE gels, and transferred to polyvinylidene fluoride membranes. For the protein detection, anti-Flag (1:5000; abmart, M20026) were used to detect Flag-CO proteins. Anti-Actin (1:5000; CWbiotech, CW0264) was used as a loading control.

### *In vitro* phosphorylation assay

For Phosphorylation assay, 1 μg BIN2-MBP/MBP and 1 μg CO-GST/mutated CO fusion proteins were added into kinase reaction buffer (25 mM Tris at pH 7.4, 12 mM MgCl_2_, 1 mM DTT, and 1 mM ATP). Then the mixture was incubated at 37°C for 1 h and boiled with 5×SDS loading buffer. The boiled reaction products were separated by SDS-PAGE with or without 50 μM Phos-Tag (NARD, AAL-107). The phosphorylated CO was detected with anti-GST antibody.

### Subcellular localization assays

The coding sequences of CO, CO^T280A^ and Q were cloned into the *pEarly-101* vector to generate YFP-CO, YFP-CO^T280A^ and mCherry-Q constructs, respectively. Then the constructs were transformed into the *Agrobacterium* strain *GV3101* and infiltrated into *N. benthamiana* leaves. The fluorescence signal of yellow fluorescent protein (YFP) was observed at 48 h post-infiltration.

## Results

### BIN2 physically interacts with CO

CONSTANS (CO) is a central regulator of floral initiation in response to photoperiod. In order to explore the regulatory effects of phosphorylation and specific protein kinases for CO, we tested a series of kinases for CO interaction through the luciferase (LUC) complementation imaging (LCI) assay. In the LCI assay, CO was fused with the amino-terminal part of LUC (nLUC) and kinases (BIN2, SK12, SnRK2.6 and MPK6) were ligated to the carboxyl-terminal part of LUC (cLUC) to generate nLUC-CO and cLUC-BIN2 (SK12, SnRK2.6, MPK6), respectively. The results showed that strong LUC activity could be observed when nLUC-CO/cLUC-BIN2 and nLUC-CO/cLUC-SK12 were co-expressed in *Nicotiana benthamiana* leaves, but no LUC activity was detected in nLUC-CO/cLUC-SnRK2.6, nLUC-CO/cLUC-MPK6 (Fig. 1a). Since BIN2 and SK12 (a homolog of BIN2) interact with CO, we further tested the interaction between BIN2 and CO through other approaches. In the yeast two-hybrid (Y2H) assay, BIN2 was fused with *pGBKT7* and CO was ligated to *pGADT7* to generate BD-BIN2 and AD-CO, respectively. The *AH109* strain yeast cells transformed with BD-BIN2 and AD-CO could grow well in the selection medium, but the negative controls did not (Fig. 1b), indicating that BIN2 physically interacts with CO in yeast cells. Consistently, the MBP-BIN2 fusion proteins could retain GST-CO proteins, whereas MBP alone could not, indicating that BIN2 directly interacts with CO in the *in vitro* pull-down assay (Fig. 1c). Moreover, we generated the *35S:Flag-BIN2/35S:YFP-CO* double transgenic plant through genetic crossing between the *35S:Flag-BIN2* and *35S:YFP-CO* lines for the Co-IP assay. The results showed that the Flag-BIN2 proteins could be precipitated by YFP-CO proteins in the double transgenic *35S:Flag-BIN2/35S:YFP-CO* seedlings in the Co-IP assay, further suggesting that BIN2 interacts with CO *in vivo* (Fig. 1d).

**Fig. 1.**
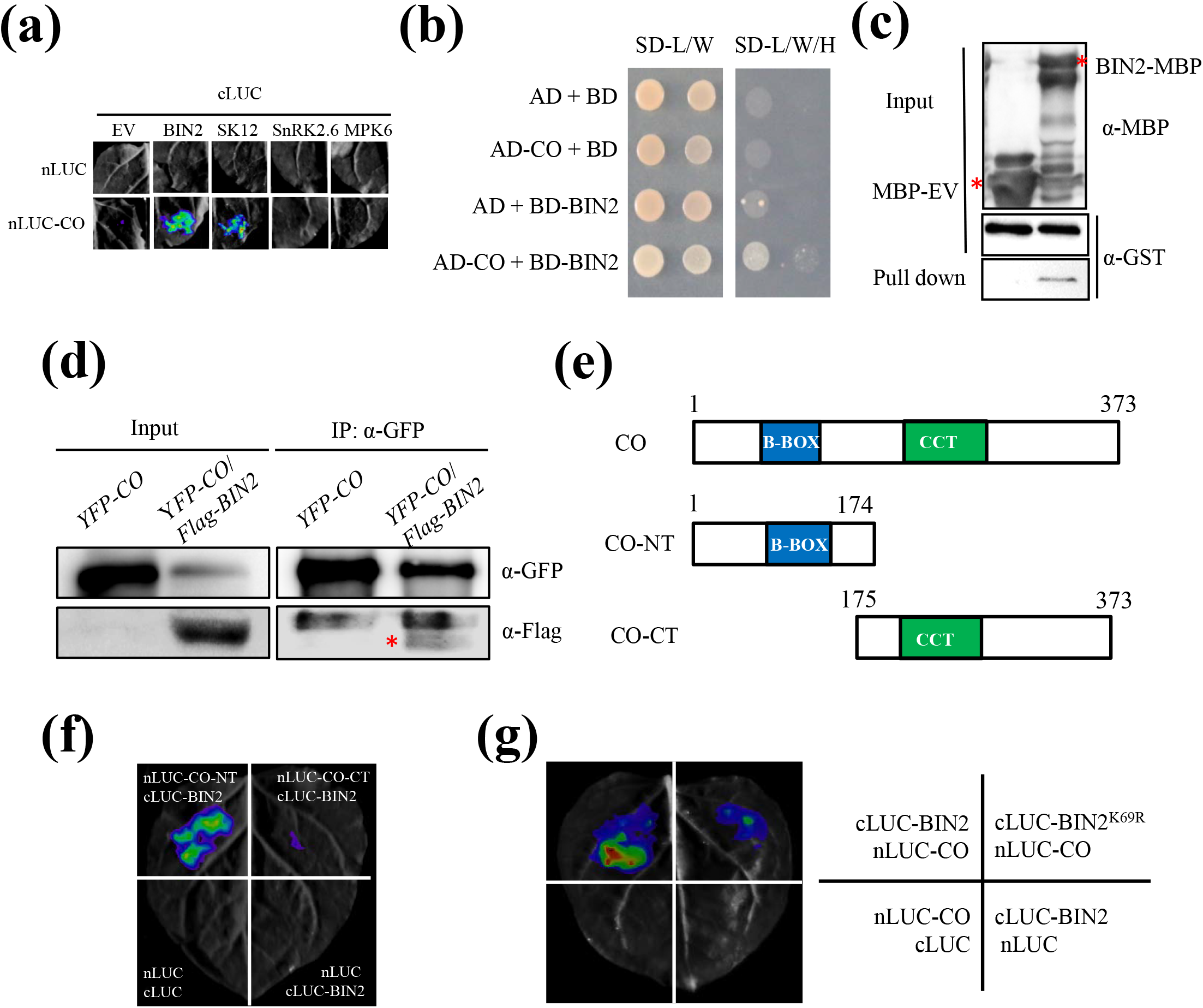
BIN2 physically interacts with CO *in vitro* and *in vivo*. (a) Identification of CO-interacting protein kinases. LCI assays showing the interaction between CO and protein kinases in *N. benthamiana* leaves. Representative images of *N. benthamiana* leaves after infiltration were shown. Empty vectors were used as negative controls. (b) Y2H assay showing the interaction between BIN2 and CO. SD-L/W, synthetic dextrose medium lacking Leu and Trp; SD-L/W/H, synthetic dextrose medium lacking Leu, Trp and His. (c) *In vitro* pull-down assay showing that BIN2 directly interacts with CO. Purified GST-CO proteins were incubated with MBP or MBP-BIN2 for the MBP pull-down assay. Red asterisk indicates specific bands. (d) Co-IP assays showing the interaction between BIN2 and CO in *Arabidopsis*. The *35S:YFP-CO* and *35S:YFP-CO/35S:Flag-BIN2* transgenic plants were grown for 10 days under LDs. Red asterisk indicates specific signal. The immunoprecipitates were detected using anti-GFP and anti-Flag antibodies, respectively. (e) Schemes displaying full-length and truncated versions of the CO protein. CO-NT, amino terminal domain of CO; CO-CT, carboxyl terminal domain of CO. (f) LCI assay showing the interaction between the truncated versions of CO and full-length BIN2 in *N. benthamiana* leaves. (g) LCI assay showing less LUC activities between BIN2^K69R^ and CO.

To map the interaction region of CO with BIN2, the full-length of CO protein was truncated into N-terminal part harboring the B-Box domain (CO-NT) and C-terminal part containing the CCT domain (CO-CT) (Fig. 1e). Those two parts of CO were fused with nLUC to generate nLUC-CO-NT and nLUC-CO-CT for LCI assays, respectively. As shown in Fig. 1f, obvious LUC activities were observed in the nLUC-CO-NT/cLUC-BIN2 co-expression samples, but only background level of LUC activities was observed in the nLUC-CO-CT/cLUC-BIN2 samples. These results suggest that the N-terminal part of CO containing the B-Box domain is required for the interaction with BIN2 (Fig. 1f). Moreover, we also explored whether the kinase activity of BIN2 is required for its interaction with CO. BIN2^K69R^ mutant form, in which the Lys69 was replaced by Arg and the GSK kinase activity was abolished, was fused with cLUC to generate cLUC-BIN2^K69R^ for LCI assays. As shown in Fig. 1g, less LUC activities were detected in the cLUC-BIN2^K69R^/nLUC-CO co-expression samples than those in the cLUC-BIN2/nLUC-CO co-expression samples (Fig. 1g), suggesting that the kinase activity of BIN2 is essential for its interaction with CO.

Since BIN2 physically interacts with CO, we were curious whether these two genes have similar expression patterns during photoperiod. To address this question, we analyzed the expression patterns of *BIN2* and *CO* in Col-0 during a 24-h photoperiod. Interestingly, we found that the transcripts of *BIN2* exhibited similar diurnal expression patterns with *CO* (Fig. S1), supporting the notion that BIN2 is supposed to post-transcriptionally modify CO protein in regulating flowering time.

### BIN2 regulates flowering time genetically upstream of CO

Although it has been known that BRs play an important role in promoting flowering in *Arabidopsis*, the underlying mechanism remains largely unclear. To explore the biological role of BR pathway in regulating flowering time, we systematically examined the flowering time phenotypes of BR biosynthetic and signaling mutants including *bin2-1, det2-9* and *bri1-116* under LD (16 h light/8 h dark) conditions. The results showed that *bin2-1, det2-9* and *bri1-116* displayed obvious late-flowering phenotypes compared with the wild type (WT) Columbia-0 (Col-0) under LD conditions (Fig. 2a, b; Fig. S2a, b). To further dissect the molecular effect contributing to the late flowering of BR biosynthetic or signaling mutants, we analyzed daily expression patterns of *FT*, a key integrator of flowering time (Putterill et al., 1995; Kobayashi et al., 1999), in these mutants using quantitative RT-PCR (qRT-PCR). The qRT-PCR assays revealed that the expression levels of *FT* in Col-0 plants were peaked at Zeitgeber time 16 (ZT16), consistent with previous report (Turck et al., 2008), but the peak of *FT* expression levels were largely abolished in the *bin2-1, det2-9* and *bri1-116* mutants (Fig. 2c; Fig. S2c). These results imply that the late flowering phenotypes of these mutants were likely due to the down-regulation of *FT* transcription.

**Fig. 2.**
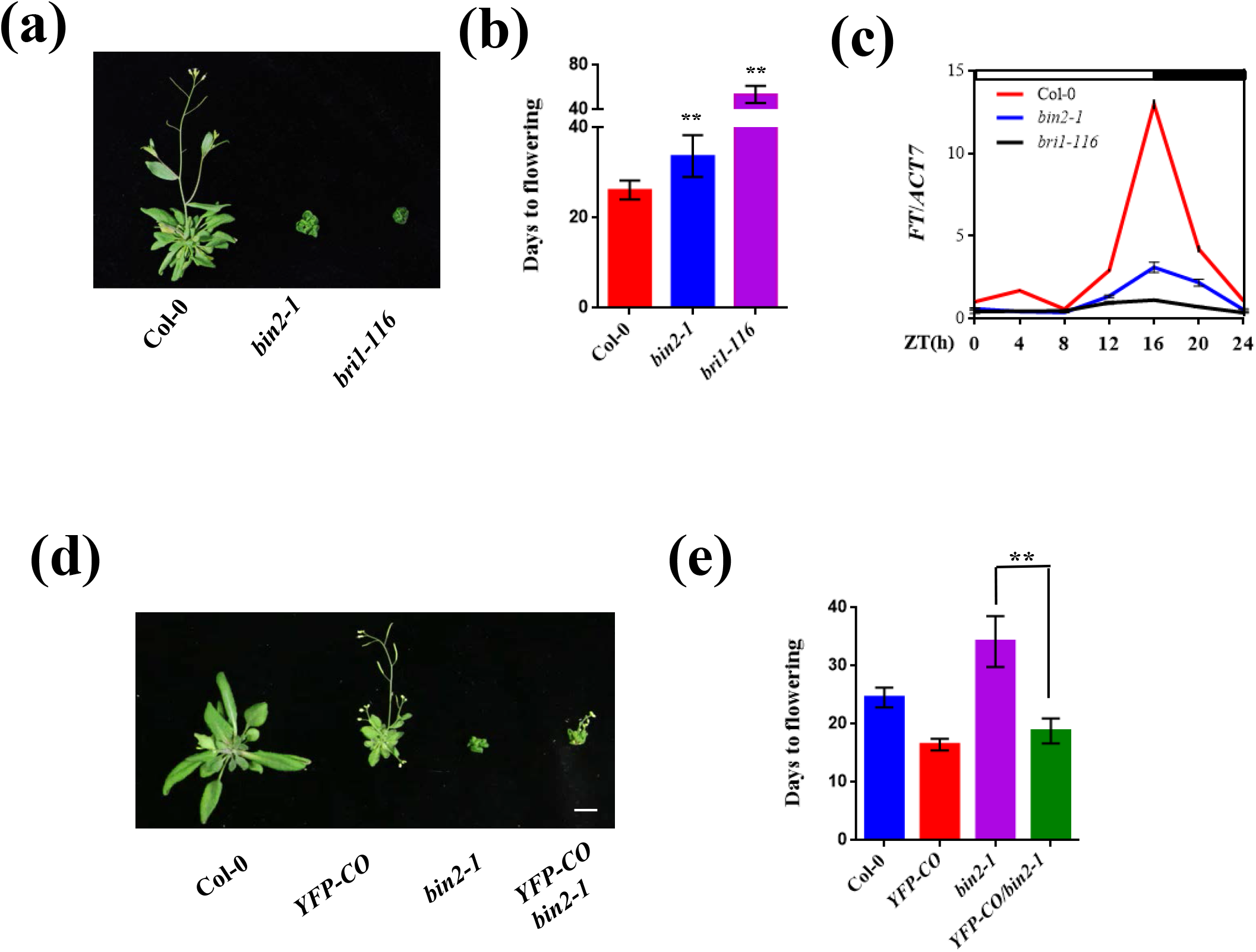
BR negatively regulates flowering time genetically upstream of CO. (a) Flowering phenotype of the indicated plants under LDs. Bar, 1 cm. (b) Days from germination to flowering of plants shown in (a). Asterisks indicate a significant difference according to Student’s *t* test (**, *P* <0.01), n > 20. (c) Diurnal expression pattern of *FT* in the indicated plants under LDs for 14 d. The mean values in Col-0 at Zeitgeber time (ZT) 0 were set to 1 (mean ± SD). White bars indicate day, and black bars represent night. (d) and (e) Genetic interaction assays between BIN2 and CO. Flowering phenotype of the indicated plants under LDs (d). Days from germination to flowering of indicated plants (e). Asterisks indicate a significant difference according to Student’s *t* test (**, *P* <0.01), n > 20. Bar, 1 cm.

Considering that CO directly activates *FT* transcription (Onouchi et al., 2000; Samach et al., 2000), we were curious about the relationship between the BIN2 and CO in regulating flowering time. To this end, we studied the genetic interaction between BIN2 and CO. We generated the *CO*-overexpressing transgenic line *35S:YFP-CO* and introduced the *35S:YFP-CO* transgene into the *bin2-1* mutant background to generate the *35S:YFP-CO/bin2-1* lines. As shown in Fig. 1d and 1e, the *35S:YFP-CO* transgenic line flowered much earlier than the wild type plant Col-0. Notably, the *35S:YFP-CO/bin2-1* line displayed obvious early-flowering phenotype compared with the *bin2-1* mutant, indicating that over expression of *CO* can rescue the late-flowering phenotype of *bin2-1* (Fig. 2d, e). This result demonstrates that BIN2 acts genetically upstream of CO in regulating flowering time.

### BIN2 phosphorylates CO at the Thr280 residue

Since the protein kinase BIN2 interacts with CO, it is important to test whether CO is a substrate of BIN2. To this end, we used a phos-tag approach, where phosphorylated proteins in the gel containing phos-tag reagent are visualized as bands with slower migration compared with the corresponding dephosphorylated proteins. As shown in Fig. 3a, the slower migrated band of CO could be observed when incubated with MBP-BIN2 and ATP; whereas this phenomenon could not be observed when incubated MBP-BIN2 without ATP (Fig. 3a). These results demonstrate that BIN2 can phosphorylate CO. Consistently, the slower migrated band of CO was not observed when incubated with MBP-BIN2^K69R^ (the kinase dead version of BIN2) and ATP (Fig. 3a), suggesting that the kinase activity of BIN2 is required for the phosphorylation of CO.

**Fig. 3.**
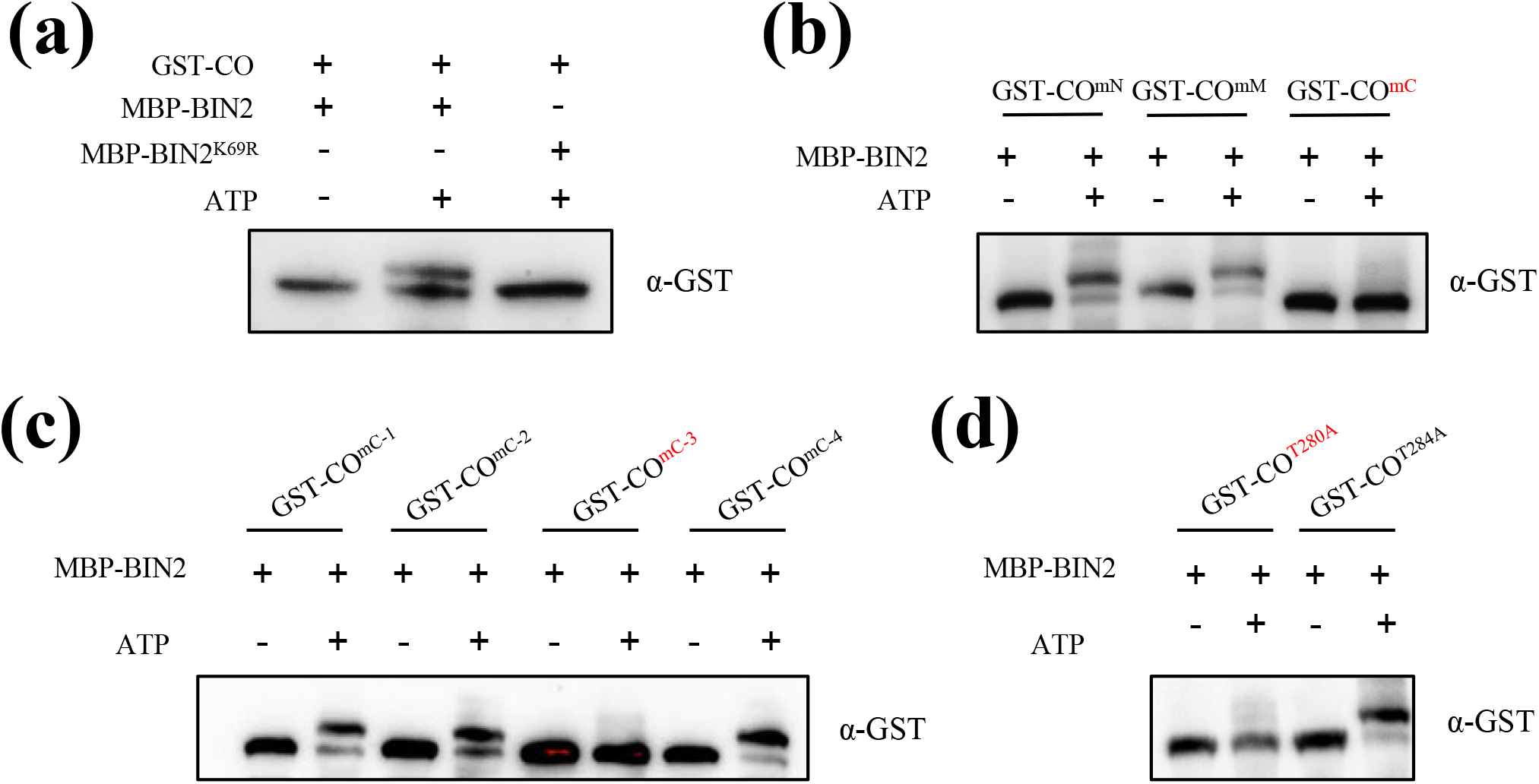
BIN2 phosphorylates CO. (a) Phosphorylation assays showing the BIN2-mediated phosphorylation of CO *in vitro*. Proteins were detected by immunoblotting with anti-GST antibody. (b) and (c) Phosphorylation assays showing the BIN2-mediated phosphorylation of CO with different mutations *in vitro*. Proteins were detected by immunoblotting with anti-GST antibody. (d) Phosphorylation assays showing the BIN2-mediated phosphorylation of CO^T280A^ and CO^T284A^. Proteins were detected by immunoblotting with anti-GST antibody.

In order to search for the putative phosphorylation sites of CO by BIN2, we first analyzed the putative BIN2 phosphorylation motifs in CO (Ser/Thr-X-X-X-Ser/Thr; Wang et al., 2002; Ryu et al., 2007, 2010). As a result, we identified 13 putative phosphorylation motifs in CO protein (Fig. S3). Next, we mutated Ser/Thr to Ala in the 1^th^-3^th^ phosphorylation motifs of CO (CO^mN^), 4^th^-9^th^ phosphorylation motifs of CO (CO^mM^) and 10^th^-13^th^ phosphorylation motifs of CO (CO^mC^) to mimic non-phosphorylated forms of CO, respectively. As shown in Fig. 3b, CO^mN^ and CO^mM^ was still phosphorylated by BIN2; in contrast, CO^mC^ was not phosphorylated by BIN2, illustrating that the phosphorylation sites are located in the 10^th^-13^th^ phosphorylation motifs of CO (Fig. 3b). Further, we mutated Ser/Thr to Ala in the 10^th^ phosphorylation motifs of CO (CO^mC-1^), 11^th^ phosphorylation motifs of CO (CO^mC-2^), 12^th^ phosphorylation motifs of CO (CO^mC-3^) and 13^th^ phosphorylation motifs of CO (CO^mC-4^), respectively. The results showed that the CO^mC-1^, CO^mC-2^ and CO^mC-4^ mutant forms were still phosphorylated by BIN2; whereas CO^mC-3^ could not be phosphorylated, indicating that the phosphorylation sites are located at the 12^th^ phosphorylation motif of CO (Fig. 3c). Finally, we mutated the Thr280 and Thr284 residues at the 12^th^ phosphorylation motif to Ala to generate the CO^T280A^ and CO^T284A^ mutant forms, respectively. As shown in Fig. 3d, CO^T284A^, but not CO^T280A^, could be phosphorylated by BIN2, suggesting that the Thr280 residue is the specific BIN2 phosphorylation site of CO protein (Fig. 3d). In summary, our efforts reveal that BIN2 can phosphorylate the Thr280 residue of CO protein.

### Phosphorylation by BIN2 represses the function of CO in promoting flowering

To explore the biological relevance of CO phosphorylation by BIN2, we generated the *pCO:Flag-CO/co-9* and *pCO:Flag-CO^T280A^/co-9* transgenic plants in the *co-9* mutant background. As shown in Fig. 4a, the *pCO:Flag-CO* transgene could completely rescue the late-flowering phenotype of *co-9*. Notably, the *pCO:Flag-CO^T280A^/co-9* lines flowered obviously earlier compared to *pCO:Flag-CO/co-9* and Col-0 (Fig. 4a–4c). The transcript levels of *CO* in these transgenic lines were comparable (Fig. 4d). Consistent with the flowering phenotypes, the transcript levels of *FT* at ZT16 in the *pCO:Flag-CO^T280A^/co-9* plants were higher than those in the *pCO:Flag-CO/co-9* and Col-0 plants (Fig. 4e). However, the CO protein levels were largely similar between the *pCO:Flag-CO^T280A^/co-9* and *pCO:Flag-CO/co-9* transgenic plants (Fig. 4f), illustrating that phosphorylation of CO by BIN2 does not influence CO protein stability. Next, we investigated whether the phosphorylation by BIN2 affects the interaction intensity between CO and TOE1/NF-YC4. The Y2H assays showed that the mutation of the Thr280 residue to Ala did not significantly affect the interaction intensity between CO and TOE1/NF-YC4 (Fig. S4). In addition, subcellular localization assay showed that the CO-YFP and CO^T280A^-YFP proteins were both localized in the nuclei, suggesting that the phosphorylation by BIN2 does not affect the subcellular localization of CO protein (Fig. S5). Taken together, these observations demonstrate that the phosphorylation of CO by BIN2 inhibits the function of CO in promoting flowering, possibly through affecting the activity of CO.

**Fig. 4.**
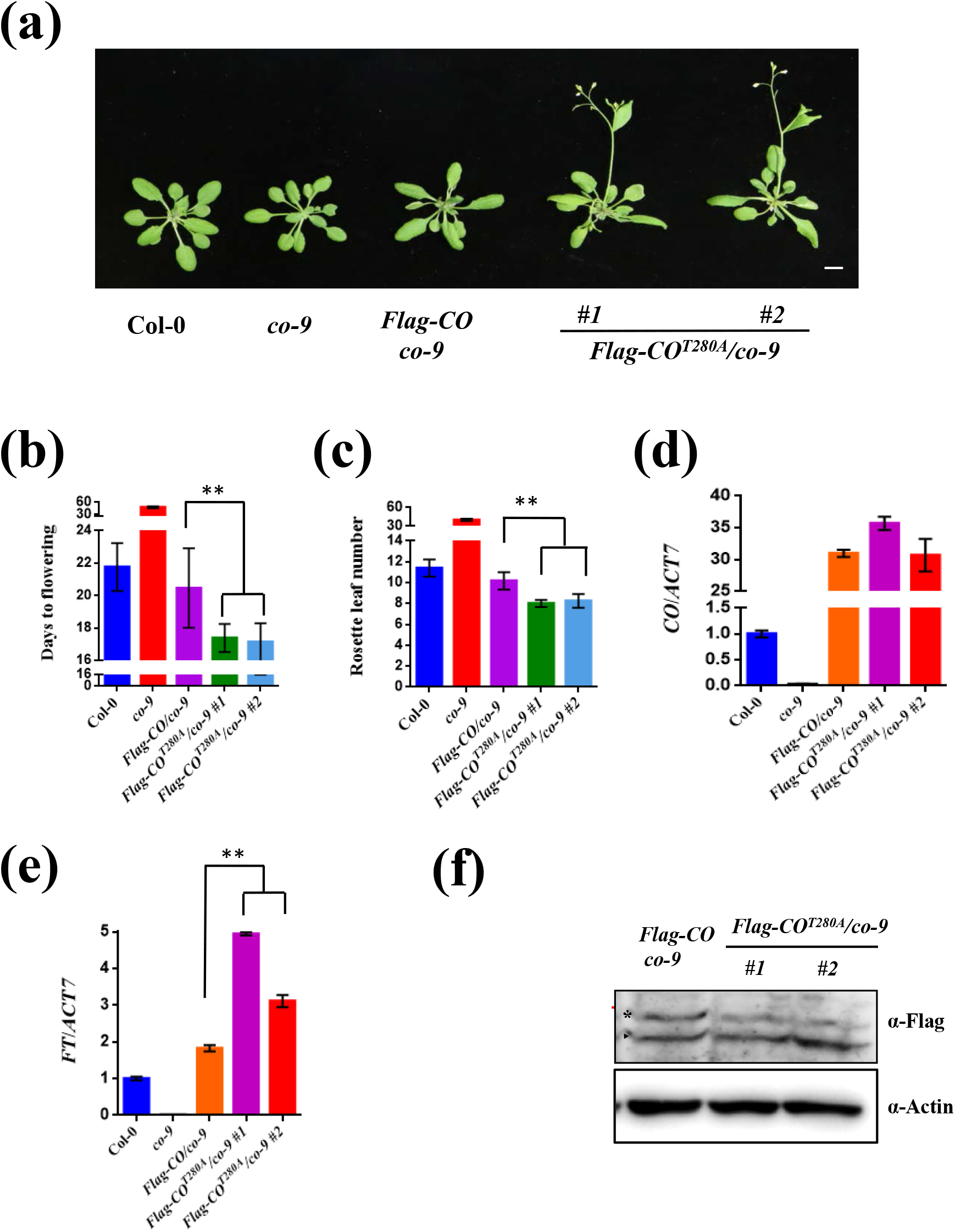
Phosphorylation of CO by BIN2 restricts its function in promoting flowering. (a) Flowering phenotype of the indicated plants under LDs. Bar, 1 cm. (b) Days from germination to flowering of plants shown in (a). Asterisks indicate a significant difference according to Student’s *t* test (**, *P* <0.01), n > 20. (c) Measurement of rosette leaf numbers of indicated plants shown in (a). Asterisks indicate a significant difference according to Student’s *t* test (**, *P* <0.01), n > 20. (d) and (e) qRT-PCR analysis showing the expression patterns of *CO* (d) and *FT* (e) at ZT16 in indicated plants under LDs. (f) The protein levels of CO at ZT16 in indicated plants under LDs. Flag-CO protein was detected by anti-Flag antibody. Actin was used as the loading control. Arrowhead and asterisk indicate unphosphorylated and phosphorylated protein forms, respectively.

### BIN2 represses the interaction of CO-CO

The formation of CO dimer/oligomer is possibly involved in influencing CO activity (Graeff et al., 2016; Lv et al., 2021). Here we showed that CO could interact with itself in plant cells (Fig. 5a), which may facilitate the formation of CO dimer/oligomer. Further, we conducted LCI assays to determine which region of CO is responsible for the interaction of CO-CO. The results showed that the N-terminal part of CO harboring the B-Box domain mediates the interaction of CO-CO (Fig. 5b). As described above, the N-terminal part of CO also mediates the interaction of BIN2 and CO (Fig. 2f). The findings that the B-Box domain of CO mediates the interaction of CO-CO and BIN2-CO prompted us to determine whether BIN2 competitively interferes with the CO-CO interaction. To test this idea, we transiently co-expressed BIN2-Flag with nLUC-CO and cLUC-CO proteins in *N. benthamiana* leaves for LCI assays. The YFP protein was expressed instead of BIN2-Flag as a control. The results showed that the samples co-expressing nLUC-CO/cLUC-CO together with BIN2-Flag displayed weaker luminescence signals compared with those of samples co-expressing nLUC-CO/cLUC-CO and YFP (Fig. 5c–5g), demonstrating that BIN2 competitively inhibits the interaction of CO-CO in plant cells.

**Fig. 5.**
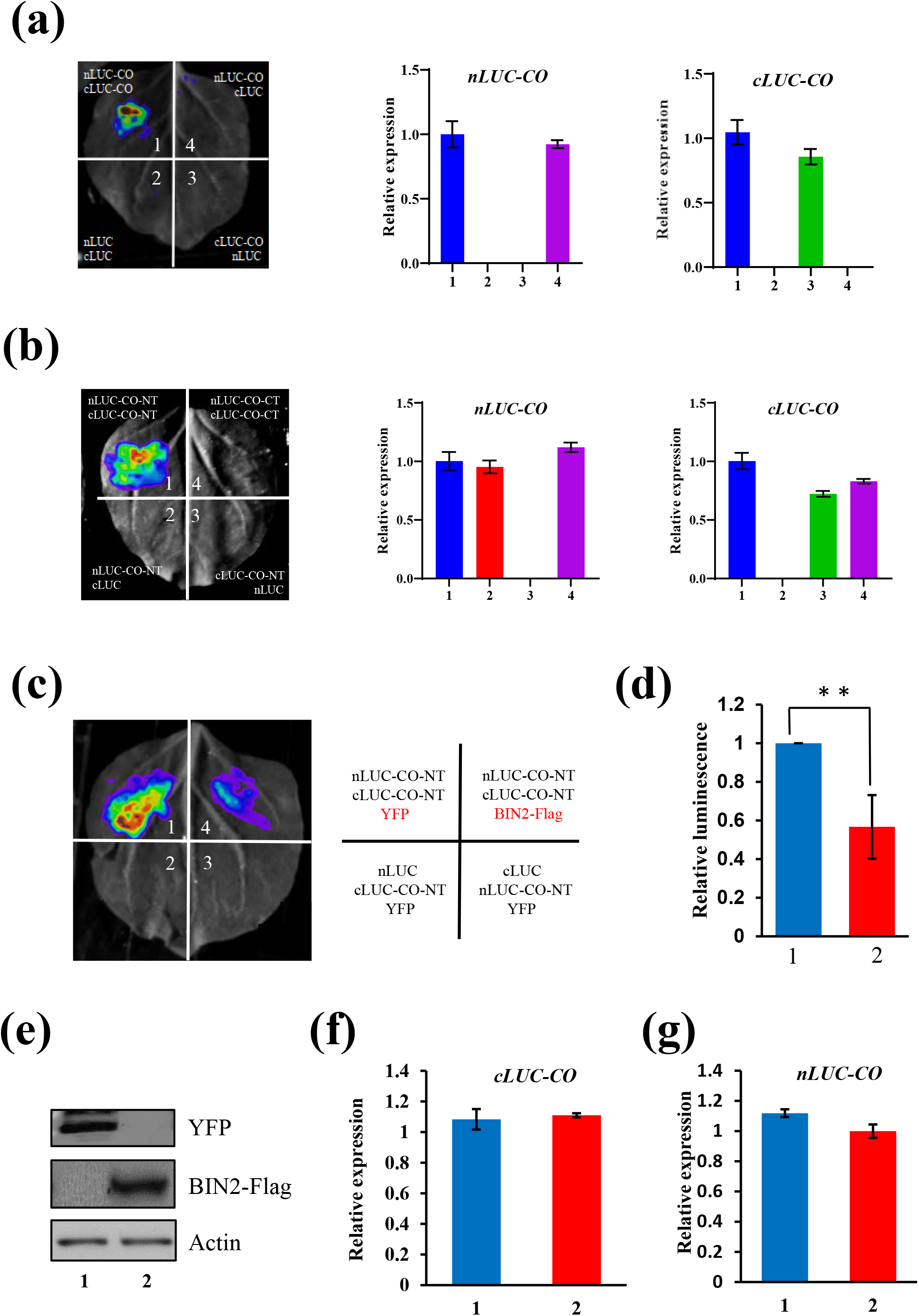
BIN2 represses the CO-CO interaction. (a) LCI assays showing the interaction of CO-CO. The expression levels of *nLUC-CO* and *cLUC-CO* in the infiltrated *N. benthamiana* leaf areas were determined by qRT-PCR. Results were normalized to *NbACTIN1* (*NbACT1*). (b) LCI assays showing that the N-terminal part of CO mediates the interaction of CO-CO. d)The expression levels of *nLUC-CO* and *cLUC-CO* in the infiltrated *N. benthamiana* leaf areas were determined by qRT-PCR. Results were normalized to *NbACTIN1* (*NbACT1*). (c) and (d) LCI assays showing that BIN2 could inhibit the interaction of CO-CO. YFP was used as a control instead of BIN2-Flag. A representative image was shown in (c), and the quantification of relative luminescence intensities was conducted in (d). Data are means ± SD. Asterisks indicate a significant difference according to Student’s *t* test (**, P <0.01). (e) The protein levels of YFP and BIN2-Flag in the infiltrated *N. benthamiana* leaf areas (infiltrations 1 and 4) were determined by immunoblotting. Actin was used as a loading control. (f) and (g) The transcript levels of *nLUC-CO* (f) and *cLUC-CO* (g) in the infiltrated *N. benthamiana* leaf areas (infiltrations 1 and 2) were quantified by qRT-PCR (means ± SD). Results were normalized to *NbACTIN1*.

## Discussion

The protein kinase BIN2 is involved in several physiological processes by phosphorylating different transcription factors. For example, BIN2 phosphorylates BZR1/BES1 to promote their protein degradation and consequently repress BR signaling; BIN2 phosphorylates ICE1 to promote its degradation in response to cold stress; BIN2 phosphorylates and stabilizes RD26 and TINY to promote drought stress response; BIN2 phosphorylates and stabilizes ABI5 to promote ABA response; BIN2 phosphorylates and inhibits SOS2 activity in regulating the salt stress response and growth recovery (He et al., 2002; Yin et al., 2002; Hu and Yu, 2014; Jiang et al., 2019; Xie et al., 2019; Ye et al., 2019; Li et al., 2020). On the other hand, a previous study reported that CO can be phosphorylated *in planta* (Sarid-Krebs et al., 2015). In this study, the phosphorylated band of CO protein can be clearly visualized; illustrating that CO can be phosphorylated by specific protein kinases. Here, we found that BIN2 physically interacts with CO *in vitro* and *in vivo* (Fig. 1a-d). And the *bin2-1* mutant displayed obvious late-flowering phenotype and overexpression of *CO* could rescue the late-flowering phenotype of *bin2-1* (Fig. 2d-e), suggesting that CO acts genetically downstream of BIN2. Next, we demonstrate that BIN2 phosphorylates the Thr280 residue of CO and restricts the function of CO in promoting flowering (Fig. 4a-f). A recent study reported that SK12 negatively regulates flowering through interacting with and phosphorylating CO at threonine 119, thus facilitating CO degradation (Chen et al., 2020). These observations suggested that the different GSK3 protein kinases BIN2 and SK12 phosphorylate the Thr280 and Thr119 residues of CO, respectively, to restrict the its function.

In a recent study, structural analysis suggested that CO might form a homomultimeric assembly via its N-terminal B-Box domain and simultaneously occupy multiple cis-elements within the *FT* promoter (Lv et al., 2021). Here, we demonstrate that the N-terminal B-Box domain of CO is responsible for the interaction of both CO-CO and BIN2-CO (Fig. 2f and 5b). Furthermore, we revealed that BIN2 competitively interferes with the CO-CO interaction that may facilitate the formation of CO dimer/oligomer (Fig. 5c, d). Thereby, we propose that BIN2 phosphorylates CO and prevents the interaction of CO-CO that might facilitate the formation of CO dimer/oligomer to restrict its function, leading to down-regulation of *FT* transcription and late flowering.

## Accession Numbers

Sequence data in *Arabidopsis* from this study can be found in the *Arabidopsis* Genome Initiative database under the following accession numbers: *BRI1* (AT4G39400), *DET2* (AT2G38050), *BIN2* (AT4G18710), *CO* (AT5G15840), *FT* (AT1G65480) and *ACTIN7* (AT5G09810), *SK12*(AT3G05840), *SnRK2.6*(AT4G33950), *MPK6*(AT2G43790).

## Acknowledgements

We thank George Coupland, Yongfu Fu, Ming-Yi Bai, Jianming Li and Zhaojun Ding for kindly providing genetic materials. This research was supported by the Central Public-interest Scientific Institution Basal Research Fund (S2022ZD02).

## Author contributions

J.S. designed this research; L.J., H.D., R.Y., Y.J. and Y.Zhang performed the experiments; J.S. and L.J. analyzed the data and wrote the manuscript; Y.Zhu and L.L. analyzed the data and revised the manuscript; K.C. provided support.

## Data availability statement

All data supporting the findings of this study are available within the paper and within its supplementary materials published online.

